# Uncertainty-Aware Tau Detection in Progressive Supranuclear Palsy Using Object Detection Models

**DOI:** 10.1101/2025.10.09.681479

**Authors:** Gracie Zhou, Tiago Azevedo, Annelies Quaegebeur, Tanrada Pansuwan, Timothy Rittman, Pietro Lió

## Abstract

Abnormal tau accumulation is a hallmark of neurodegenerative tauopathies such as Progressive Supranuclear Palsy (PSP). Traditional post-mortem assessments rely on manual lesion annotation, which is time-consuming and subjective. Existing machine learning methods typically involve multi-stage, feature-based pipelines, resulting in limited scalability and reliance on handcrafted features.

This work evaluates deep learning-based object detection (OD) as a unified, end-to-end alternative for automating tau lesion identification and classification. We fine-tuned YOLO and Faster R-CNN models on a dataset of 29,852 tau lesions from 16 PSP brain slides, achieving mAP@50 of 0.702. To address the limitations of conventional OD models, which assume overconfident spatial predictions, we integrated Monte Carlo Dropout into both architectures to estimate spatial and label uncertainty. We evaluated detection uncertainty using Probability-based Detection Quality. Our results demonstrate that incorporating uncertainty offers a more comprehensive evaluation of detection reliability, complementing traditional metrics like mAP. For clinical application, we integrate a non-stochastic YOLO model into QuPath for whole-slide inference. Downstream clinical validation was conducted, including SHAP-based saliency analysis and correlation between model-predicted tau densities and PSP Rating Scale scores, a clinical disease severity measure. The alignment of detection outputs with pathological features and clinical severity underscores the clinical utility of OD models.

Overall, this work highlights the complementary strengths of deep object detectors and uncertainty quantification, offering a path toward scalable, interpretable, and clinically meaningful tau pathology analysis. Code is available at https://github.com/Pidongg/dl_histopathology.

## I. Introduction

Tau protein is crucial for the stabilisation of cellular microtubules. Neurodegenerative diseases characterized by abnormal tau accumulation are known as tauopathies, which include Alzheimer’s Disease and Progressive Supranuclear Palsy (PSP). The distribution of abnormal tau in neurons and glial cells exhibits distinctly recognizable patterns, and correlates with clinical symptoms [1], [2]. For PSP, tau aggregates in isolation, unlike Alzheimer’s disease, where tau appears alongside beta-amyloid aggregation. Tau aggregates in PSP can be classified into four distinct types: Tufted Astrocytes (TA), Coiled Bodies (CB), Neurofibrillary Tangles (NFT), and fragments of tau, which are seen on histological examination as representing the processes of affected astrocytes, or tau released from cells having undergone apoptosis or necrosis. Post-mortem detection of tau cells in brain tissue is valuable for pathologists to identify a specific neuropathology and to understand disease mechanisms.

However, current post-mortem assessments are semi-quantitative, requiring pathologists to manually identify, quantify, and grade disease severity on a simple ordinal scale [3]. This approach is time-consuming and subject to inter-rater variability, which limits the scale of the analysis to specific brain subregions. Machine Learning (ML) enables automated tau detection, significantly reducing the time required for quantification. The increased efficiency allows researchers to identify more associations between tau distribution and tauopathies, advancing disease understanding [4].

Current ML research in tau detection can be categorized into two main groups: feature-based approaches and deep learning methods. Feature-based approaches commonly employ a two-stage process: a segmentation model extracts features as input into a separate feature-based classification model that assesses disease severity based on these features [5]. We have previously applied this methodology [4], combining QuPath’s built-in thresholder tool for initial tau object segmentation, then extracting 54 features for each segmented object to feed into random forest classifiers, to identify four distinct tau aggregates. In contrast, deep learning approaches offer a potentially more efficient end-to-end solution that integrates detection and classification into a single stage, eliminating the need for preliminary segmentation tools and feature extraction. A deep learning approach has been applied to tauopathies by Koga et al, who developed object detection (OD) models using the well-established YOLO architecture to identify and classify patterns of tau pathology [6]. The outputs were then passed to random forest classifiers to distinguish between neurodegenerative pathologies, with a promising accuracy of over 95%. However, this study did not classify individual tau aggregates, so it is not possible to establish the burden of tau neuropathology in a given individual.

We present an uncertainty-aware OD framework for histopathology, applied to tau pathology in PSP. We build on our previous work using the same dataset, and replacing the deterministic, feature-engineered pipeline with an end-to-end deep learning approach. Our aim was to design a pipeline that does not require preprocessing, integrates uncertainty estimation, and can be deployed directly within pathology workflows. Our main contributions are:

1. Uncertainty-aware OD pipeline for tau pathology - We develop and fine-tune YOLO11 and Faster R-CNN models for tau detection in PSP, incorporating a complete preprocessing pipeline and a QuPath plugin that sup-ports custom model inference and whole-slide tiling. This enables seamless integration into existing pathology workflows for efficient analysis.
2. Integration of probabilistic OD in histopathology - To the best of our knowledge, this is the first adaptation of Monte Carlo Dropout (MC-Drop) to quantify both spatial and classification uncertainty in tau detection, with evaluation via Probability-based Detection Quality (PDQ), a metric not previously applied in this domain. This allows quantifying confidence in both localisation and classification, providing pathologists with richer information than traditional metrics.
3. Deployment-oriented evaluation and interpretability framework - We generated SHAP-derived saliency maps, feature map visualisations, and analysis of correlations between model-inferred tau densities and PSP Rating Scale (PSPRS) to interpret model predictions and confirm focus on pathologically relevant features. These tools support model transparency, facilitate expert trust, and inform human-in-the-loop workflows.

## II. Dataset

The dataset used in this paper comprises slides from 32 post-mortem human brains, donated by people with PSP to the Cambridge Brain Bank, with ethical approval from the Wales Research Ethics Committee. Slides are organised by patient and corresponding brain regions for each patient. Preprocessing of 16 slides was performed in our previous study, resulting in 29,852 labelled tau instances. An additional 56 slides without tau labelling were available and used for correlation analysis between predicted tau density and PSPRS scores per patient.

Further data processing was necessary to adapt the dataset for OD models. Whole-slide images measured approximately 4.9 billion square pixels, whereas tau objects were substantially smaller, averaging under 10,000 square pixels. To facilitate detection, each slide was divided into 512 × 512 pixel tiles with a 32-pixel overlap to reduce the number of cut-off tau objects at tile boundaries. Annotations were then converted into bounding boxes in a format compatible with OD models, and tiles without any labelled tau objects were excluded. The dataset was split such that 7 slides used in our prior study were retained as a held-out test set, while the remaining 9 were allocated to training and validation. 1 validation slide was randomly selected from each brain region to ensure representation of region-specific morphological variations during model development. To improve model robustness, empty tiles were included in training and validation to penalize false positives. The number of such tiles was limited to match the number of minority class object tiles per slide to avoid class imbalance.

The distribution of the final dataset size after processing is shown in Figure 1.

**Fig. 1.**
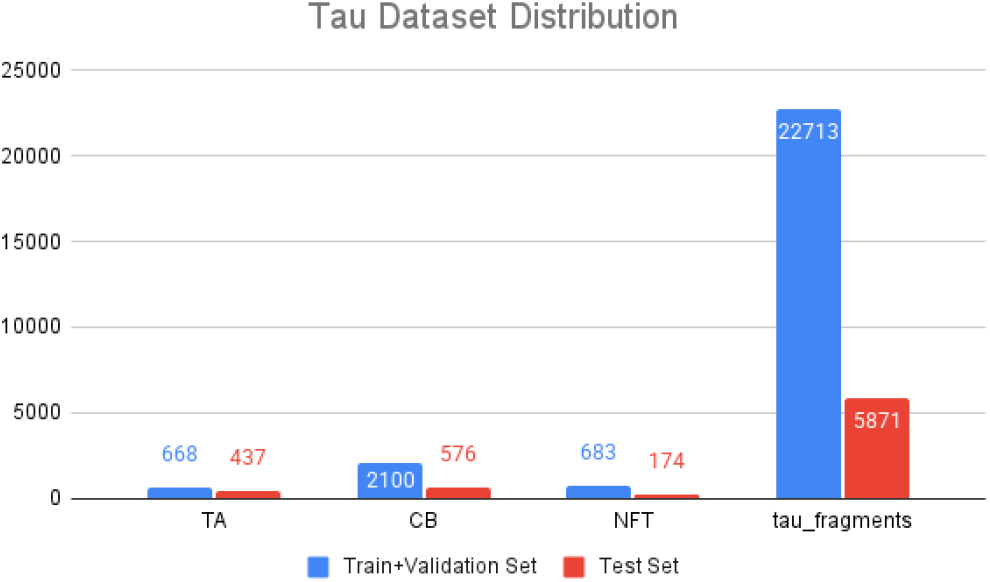
Distribution of labeled tau objects. The horizontal axis refers to the 4 types of tau present in this dataset: TA, CB, NFT, Tau Fragments.

## III. Non-stochastic model development

This work employs two open-source models: *Ultralytics*’ YOLO11 [7] and *PyTorch*’s Faster R-CNN with ResNet50 backbone [8]. Both were previously pre-trained on the COCO dataset [9], then fine-tuned on our tau dataset.

Model development involved selecting appropriate data augmentation strategies, tuning hyperparameters, and fine-tuning the models.

### A. Data Augmentation

To address dataset limitations and mitigate overfitting, data augmentation was applied during training through random transformations of input samples. Augmentation methods were selected based on two key principles: alignment with how pathologists interpret tau pathology, and preservation of object identity. Since tau identification depends primarily on shape, color, and local context, augmentations that alter color were excluded to avoid disrupting critical features. Only invariant transformations—those that preserve the fundamental identity of tau objects—were considered.

Each augmentation strategy was evaluated individually, and the best-performing ones were combined. The most effective methods included Contrast Limited Adaptive Histogram Equalization, which enhanced local contrast to simulate variations in staining, lighting, and tissue preparation; orientation-based augmentations like flipping, rotation, and translation, which preserved object structure and supported spatial invariance; morphometric transformations such as scaling and shearing, which simulated natural variation in tau morphology; and kernel-based blurring, which improved robustness to imaging noise for Faster R-CNN, though not for YOLO11.

### B. Hyperparameter Tuning

Model hyperparameters, including learning rate schedules, weight decay, optimizer settings, and loss function weights, were tuned using Ray Tune with random search [10]. During this process, an important insight emerged: tau fragment classes achieved better detection accuracy at lower confidence thresholds than other classes. Many correct detections of these fragments produced relatively low confidence scores, suggesting that a uniform threshold for Non-Maximum Suppression (NMS) was suboptimal. Consequently, the models were improved by introducing class-specific confidence thresholds for NMS. Final model selection for both YOLO11 and Faster R-CNN was based on mAP@50 performance on the validation set. Various combinations of per-class confidence and IoU thresholds were tested, and the settings that yielded the highest mAP@50 on the validation set were chosen as optimal.

### C. Traditional metrics results

The optimized YOLO11 and Faster R-CNN models are evaluated using traditional metrics—mAP@50, F1-score—and compared against benchmarks from the feature-based pipeline in our previous study [4]. Overall, YOLO11 slightly outperformed Faster R-CNN on the test set for all metrics (Figure 2). Among the four types of tau, TA and NFT achieved the highest detection mAP@50, followed by CB (Figure 3). This is consistent with prior findings—for example, a YOLOv3-based study reported 0.63 for CB and 0.85 for TA [6], underscoring CB’s detection challenges. Tau fragments classification proved particularly difficult, partially due to their inherent morphological variability. As detached portions of larger tau objects, fragments exhibit diverse shapes and sizes, complicating both detection and classification.

**Fig. 2.**
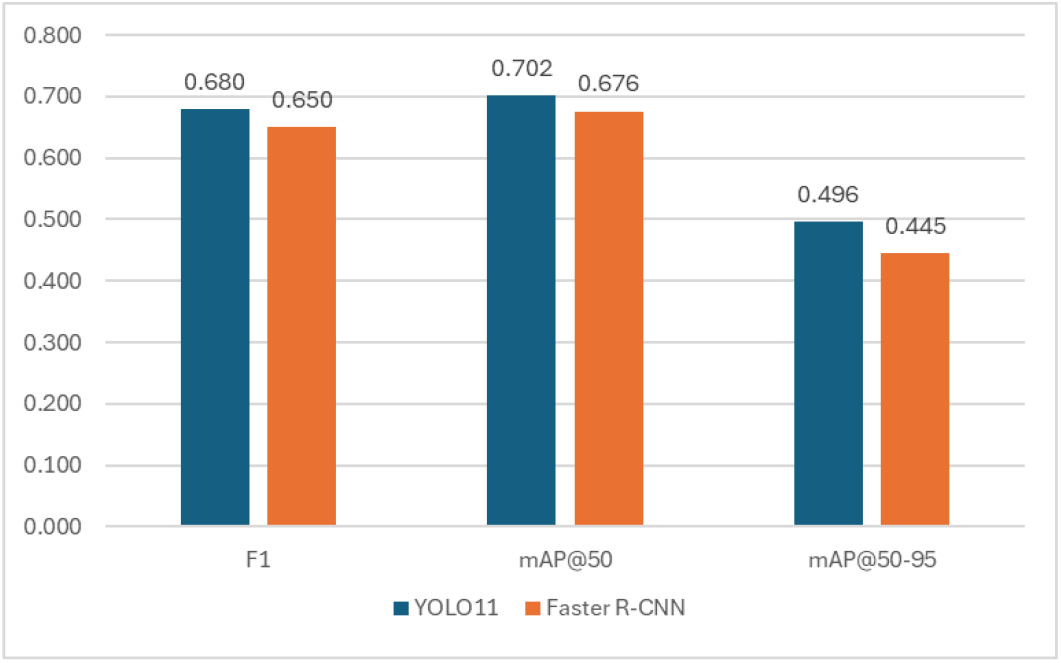
Comparison of YOLO11 and Faster R-CNN model performance on the held-out test set. Metrics shown are mAP@50 and F1-score per tau type. YOLO11 slightly outperforms Faster R-CNN overall, with the highest accuracy for TA and NFT, and lowest for tau fragments.

**Fig. 3.**
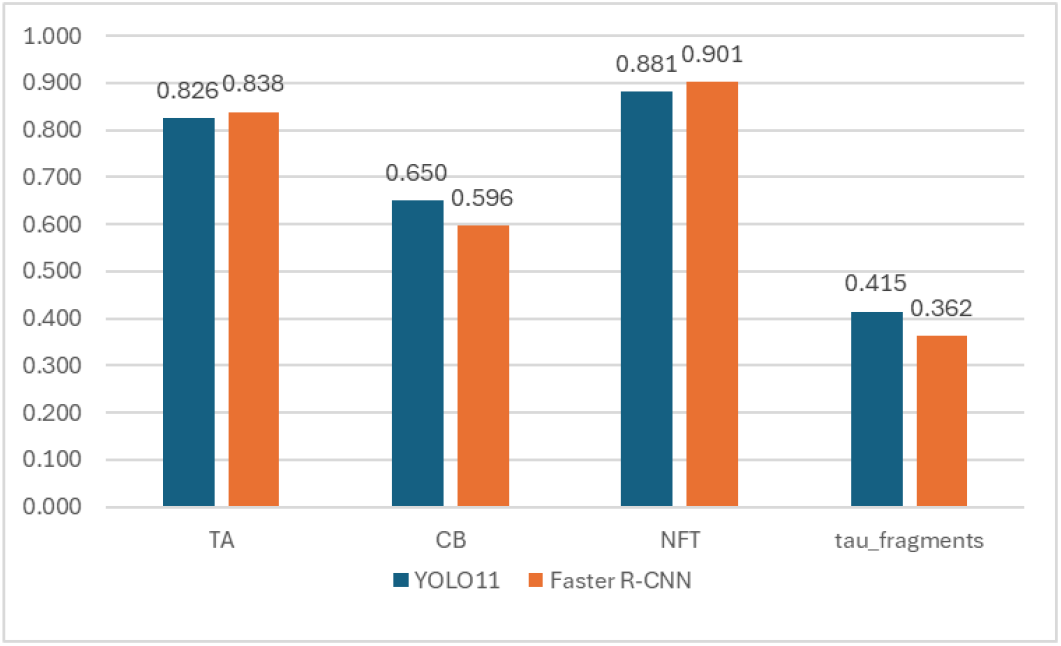
mAP@50 for each tau class using the best YOLO11 and Faster R-CNN models. TA and NFT achieve the highest scores, CB shows intermediate performance, and tau fragments remain the most challenging class for accurate detection.

#### 1) Comparison with Feature-based Approach’s Results

Before comparing the feature-based and OD approaches, it is important to note a key evaluation discrepancy. Our previous work [4] used random forest classifiers on objects detected by QuPath’s thresholder, with ground truth annotations limited to these detections. In contrast, our OD models operate directly on raw images, enabling detection of tau objects missed by the thresholder and thus absent from the ground truth. This creates an asymmetric evaluation: correctly detected but unannotated tau objects are penalized as false positives. The issue is especially pronounced for tau fragments, which are abundant and only partially annotated due to practical constraints—a common challenge in histopathology, where complete annotation is rarely feasible. Consequently, training data quality is suboptimal and evaluation metrics may underestimate true model performance.

This partially explains the disparity in tau fragment performance. While the feature-based approach reported 99.49% recall for tau fragments, the OD model showed lower recall and mAP@50, with fragments frequently confused with background and not misclassified as other tau types (see Figure 4).

**Fig. 4.**
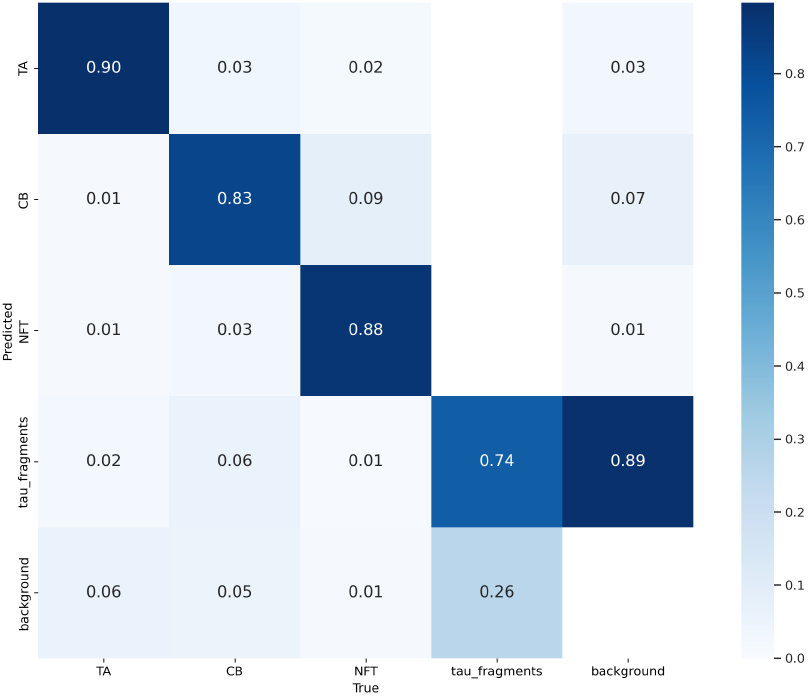
Confusion matrix for the best YOLO11 model, showing true and predicted class labels for the test set.

Despite these differences, the feature-based results offer valuable benchmarks. That method achieved F1 scores exceeding 0.9 for most classes across all brain regions. The OD model’s lower overall F1 score of 0.68 reflects evaluation under a more challenging framework, yet still demonstrates promise for deep learning in histopathology—particularly as more fully annotated datasets become available.

Importantly, our findings support the use of OD models within a ‘human-in-the-loop’ annotation framework. By detecting tau fragments overlooked by thresholding, OD can accelerate and enhance expert annotation, directly addressing the limitations that currently affect evaluation and training quality.

#### 2) Per-brain-region Results Comparison

Given morphological variability across regions, our previous work trained separate random forest classifiers for cortical, dentate nucleus (DN), and three basal ganglia (BG) re-gions: putamen, globus pallidus (GP), and subthalamic nucleus (STN) [4]. Similarly, we trained YOLO11 and Faster R-CNN models per region, using only data from the corresponding region. Due to limited data, the three BG regions were grouped for OD training. We also trained general models on the full multi-region dataset. All models were evaluated on region-specific test subsets. Contrary to expectations, general models consistently outperformed region-specific ones—likely benefiting from transfer learning and larger training sets.

Feature-based results were best in putamen, followed by cortical, GP, DN, and STN. OD models showed a similar trend: highest performance in cortical, then BG, with DN lowest—partly due to the absence of TA, a class generally easier to detect. Both approaches struggled with CB in cortical and DN, often misclassified as tau fragments. OD models, for instance, misclassified 25% of CB as tau fragments in DN (Figure 4). This alignment in performance patterns across methods suggests OD models effectively capture the morphological features that drive classification in feature-engineered approaches, without the need for manual thresholding.

### D. QuPath Plugin Development

We integrated inference using our non-stochastic YOLO11 model into QuPath using the *qupath-extension-djl* plugin, built on the Deep Java Library (DJL)—a high-level framework supporting multiple deep learning engines [11], [12]. This integration enhances QuPath’s utility for histopathological image analysis, making deep learning-based tools accessible to pathologists.

While promising, the existing plugin is still under development and lacks several core features essential for histopathology workflows. First, it currently supports only models from DJL’s model zoo, which includes general-purpose architectures trained on datasets like COCO [9]. These are not tailored to histopathological tasks, where detecting disease-specific morphological features requires custom-trained models. Second, due to the scale difference between whole-slide images and the small size of tau objects, automatic tiling is necessary to ensure comprehensive analysis—yet this functionality is not natively supported. Finally, in practical applications, inference is often required only within specific anatomical regions, but region-of-interest (RoI) selection is not available in the standard plugin.

To address these limitations, we extended the plugin to support custom models, implement automated tiling, and enable RoI-based inference. These additions make the plugin more suitable for real-world histopathology applications, supporting efficient and targeted analysis.

## IV. Uncertainty integration

Knowing the uncertainty of each detection is crucial for pathologists to distinguish between inherently unclassifiable tau objects and to have a more comprehensive view of each detection. In this work, uncertainty is quantified using MC-Drop and evaluated using the PDQ metric, which assesses both spatial and label uncertainty quantification in detections.

### A. MC-Drop

MC-Drop provides a practical approximation to Bayesian inference by enabling stochastic inference in deterministic neural networks [13]. At inference time, dropout layers are kept active, and multiple forward passes are performed on the same input. This produces a distribution of detection outputs, which captures uncertainty—higher variance in outputs for a given detection corresponds to greater uncertainty. For each detected object across these passes, two distributions are obtained:

- **Bounding box coordinates**: Capture spatial uncertainty.
- **Class probabilities**: Capture label uncertainty.

While the literature shows no consensus on optimal dropout placement for MC-Drop, our implementation prioritized effective uncertainty estimation while minimizing computational overhead and preserving model performance. Therefore, dropout layers were inserted into the detection heads of YOLO11 and Faster R-CNN, affecting only a small subset of model weights. This strategic placement enables a potential computationally efficient inference process: during inference, multiple forward passes can be performed through only the final network layers while keeping earlier layers’ outputs cached.

For YOLO11, two dropout layers were added before the last C3K2 block for each detection head. For Faster R-CNN, we added one dropout layer before each of the last two convolutional layers in the box regressor part of the architecture.

Six stochastic models were trained across the two architectures, using dropout rates of 0.25, 0.5, and 0.75 to systematically assess the impact of dropout intensity on performance. All models were subsequently fine-tuned on the tau dataset using architecture-specific optimal hyperparameters determined in earlier experiments.

### B. Evaluating uncertainty with PDQ

Unlike traditional metrics focusing on binary correct/incorrect detection outcomes, PDQ rewards accurate uncertainty quantification—low uncertainty for correct detections and high uncertainty for wrong detections. It has two components:

1. **Label quality**: Captures classification confidence for the correct class.
2. **Spatial quality**: Measures how well the predicted spatial probability distribution aligns with the ground truth mask. Spatial quality reaches 1 when a detection assigns full probability to the ground truth pixels and none elsewhere.

The pairwise PDQ (pPDQ) score is the geometric mean of label and spatial quality for each matched detection–ground truth pair, with global assignment optimized per image. PDQ requires both bounding boxes and pixel-level masks, enabling evaluation of uncertainty in both classification and localization. For implementation, we first extracted segmentation masks from tiled QuPath images with visual verification tests. For label quality evaluation, the NMS functions in both YOLO11 and Faster R-CNN were modified to retain all class confidence scores instead of keeping only the highest.

For MC-Drop, we adapted *stochastic-YOLO* [14], whose open-source code in YOLOv3 using Darknet required an adaptation to our PyTorch implementation of YOLO11 and Faster R-CNN. Dropout layers were activated at inference, and ten stochastic forward passes per tile were performed. Class scores and box coordinates were averaged pre-NMS to compute final detections. NMS was applied to the mean predictions to determine final detections. For each retained detection, covariance of bounding boxes was calculated across samples and stored alongside the mean prediction.

For PDQ evaluation, we modified the open-source *pdq evaluation* repository [14] to convert our custom predictions to RVC1 format and load segmentation masks and ground truths directly from local directories.

### C. PDQ evaluation results

Overall, increasing dropout rate leads to increased spatial quality but decreased label quality and mAP scores. The full results are shown in Table I, where the increase in spatial quality is observed to be substantial—around 3 times from baseline to 0.75 dropout rate for each architecture and confidence threshold combination, confirming that MC-Drop effectively captures spatial uncertainty in covariance matrices of bounding box coordinates. The drop in label quality and mAP can be explained by dropout layers deliberately introducing variability in predictions, making class probabilities more diffuse. This behaviour was similarly seen in previous literature [14].

**TABLE I.** PDQ, spatial quality, label quality, and mAP@50–95 for YOLO11 and Faster R-CNN models at various dropout rates (0, 0.25, 0.5, 0.75) and confidence thresholds (0.1, 0.25, 0.5). The highest value for each metric is highlighted in yellow.

| Model | PDQ | Spatial Quality | Label Quality | mAP@50-95 |
| --- | --- | --- | --- | --- |
| rcnn_0.1_baseline | 0.010 | 0.056 | 0.604 | 0.433 |
| rcnn_0.1_0.25 | 0.016 | 0.104 | 0.548 | 0.437 |
| rcnn_0.1_0.5 | 0.017 | 0.111 | 0.559 | 0.435 |
| rcnn_0.1_0.75 | 0.019 | 0.131 | 0.513 | 0.349 |
| rcnn_0.25_baseline | 0.016 | 0.057 | 0.665 | 0.428 |
| rcnn_0.25_0.25 | 0.025 | 0.111 | 0.624 | 0.431 |
| rcnn_0.25_0.5 | 0.027 | 0.121 | 0.632 | 0.428 |
| rcnn_0.25_0.75 | 0.030 | 0.138 | 0.581 | 0.325 |
| rcnn_0.5_baseline | 0.023 | 0.061 | 0.744 | 0.410 |
| rcnn_0.5_0.25 | 0.039 | 0.131 | 0.719 | 0.413 |
| rcnn_0.5_0.5 | 0.043 | 0.140 | 0.719 | 0.418 |
| rcnn_0.5_0.75 | 0.043 | 0.140 | 0.664 | 0.252 |
| yolo11_0.1_baseline | 0.015 | 0.050 | 0.362 | 0.443 |
| yolo11_0.1_0.25 | 0.018 | 0.067 | 0.297 | 0.337 |
| yolo11_0.1_0.5 | 0.025 | 0.091 | 0.285 | 0.400 |
| yolo11_0.1_0.75 | 0.035 | 0.139 | 0.261 | 0.375 |
| yolo11_0.25_baseline | 0.019 | 0.060 | 0.487 | 0.397 |
| yolo11_0.25_0.25 | 0.019 | 0.086 | 0.450 | 0.300 |
| yolo11_0.25_0.5 | 0.026 | 0.120 | 0.438 | 0.356 |
| yolo11_0.25_0.75 | 0.031 | 0.184 | 0.414 | 0.332 |
| yolo11_0.5_baseline | 0.017 | 0.090 | 0.675 | 0.331 |
| yolo11_0.5_0.25 | 0.010 | 0.087 | 0.643 | 0.221 |
| yolo11_0.5_0.5 | 0.016 | 0.176 | 0.666 | 0.271 |
| yolo11_0.5_0.75 | 0.016 | 0.265 | 0.624 | 0.233 |

The Faster R-CNN model, with a confidence threshold of 0.5 and a dropout rate of 0.5, achieves the best PDQ score of 0.043. Upon investigation, Faster R-CNN tends to produce many false positives at low confidence levels, which explains the higher label quality at higher confidence thresholds. The YOLO11 model with the highest PDQ score (0.035) has a dropout rate of 0.75 and a confidence threshold of 0.1. From our results, YOLO11 models tend to have higher spatial quality than Faster R-CNN, while Faster R-CNN tends to have better label quality. This is contrary to the findings by existing research [15], which can be attributed to the architectural difference between our versions of YOLO. The highest mAP was observed in non-stochastic YOLO11, with a 0.1 confidence threshold, despite having poor PDQ, spatial, and label qualities. This is confirmed by other research [14], [15] and highlights a well-known issue with mAP: it can be artificially inflated by lowering confidence thresholds, giving a false sense of good performance, masking the presence of false positives. This occurs because predicting more boxes increases the likelihood of at least one box overlapping with a ground truth object, which might lead to matched ground truth that would otherwise be unmatched, therefore increasing precision, which tend to out-weigh the negative effect of extra bounding boxes generated for the same object (some of which are eliminated by NMS anyway). PDQ, on the other hand, penalises both false positives and false negatives, and is designed to evaluate systems for real-world deployment, where false positives could be harmful. It also evaluates spatial quality and quantifies uncertainty, which is useful for safety-critical applications like pathological imaging.

Also, notice the low spatial quality of non-stochastic models, as conventional object detectors assume full confidence in their bounding box locations, which aligns with existing research [14], [15].

## V. Detection Interpretation

Despite strong performance, OD models remain difficult to trust in clinical settings due to their black-box nature. To improve interpretability, we used SHAP [16] to generate saliency maps, which visualize how different image regions contribute to detection confidence (Figure 5).

**Fig. 5.**
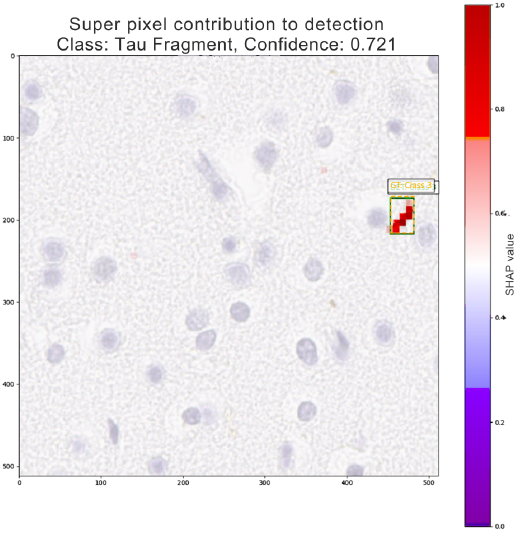
Saliency map for a detected tau fragment. Red regions contribute positively to detection confidence; blue regions contribute negatively.

As SHAP was originally designed for models with discrete input features, we adapted an existing implementation [17] for OD that uses superpixels to approximate feature groups. Each superpixel was iteratively masked, and the resulting change in detection quality was measured using a composite score, defined as the product of confidence and IoU for the best-matching detection of the correct class. Weighted linear regression was then used to estimate each superpixel’s contribution, with hue and saturation in the resulting saliency map indicating direction and magnitude of impact. Hyperparameters such as superpixel size and number of regression samples were tuned by were tuned empirically to optimize interpretability while minimizing noise and runtime.

Using this adapted approach, SHAP analysis was applied to representative examples across tau classes and brain regions using the best-performing YOLO11 model. Regions with darker staining consistently received higher SHAP values, indicating that stain intensity is a key discriminative feature. Additionally, high-contribution regions often formed class-specific morphologies—such as star-like patterns for TA, strands for tau fragments, and dense central regions for NFT—suggesting the model relies on pathologically meaningful features in its classification decisions.

These findings provide another (albeit more subjective) evaluation layer complementing the previously established evaluation framework, demonstrating that the model focuses on pathologically relevant features.

## VI. Correlation with PSPRS Scores

To evaluate the clinical relevance of model-predicted tau densities, we analysed their association with Progressive Supranuclear Palsy Rating Scale (PSPRS) scores, a validated measure of symptom severity in PSP, where higher scores indicate greater impairment.

Tau density was estimated on a brain-region basis on subcortical slides through our QuPath plugin using the best YOLO11 model. Densities were computed as the number of tau objects per *µ*m^2^, with additional metrics including total tau density and hallmark density (comprising all tau types except fragments). Due to skewed distributions, a logarithmic transformation was applied before statistical analysis.

Spearman’s rank correlation coefficient was used to evaluate associations, with significance defined at p ¡ 0.05. A significant positive correlation was observed between PSPRS scores and both CB and hallmark tau densities within DN. In contrast, correlations in BG regions were generally weaker and non-significant. Tau fragment density showed weak negative correlations with clinical severity, potentially reflecting limitations in the model’s detection accuracy for this class. No significant correlations were found when aggregating tau densities across all subcortical regions.

These findings are consistent with previous feature-based approaches, which also reported stronger associations between tau density and clinical severity in DN compared to other regions. The alignment reinforces the potential of OD models to generate clinically meaningful pathology metrics.

## VII. Conclusion

We developed models and tools that assist pathologists in detecting tau objects in post-mortem images efficiently. This is achieved through developing state-of-the-art OD models ss(YOLO11 and Faster R-CNN), and stochastic versions that enable the quantification of detection uncertainty. The models are evaluated using both traditional and probabilistic metrics, providing a comprehensive overview of their performance. The most optimal YOLO11’s reasoning process is further interpreted through SHAP and feature map visualisations, providing insights into how the model identifies different tau pathology classes based on color, shape, and contextual features. The development of a QuPath plugin enables the execution of custom-built models on the QuPath software, which is particularly useful for pathologists as fine-tuned models often outperform pre-trained models and are commonly used for histopathological OD. Finally, correlation analysis between tau density and PSPRS is performed; the significance of specific correlations shows the promise of the OD model for understanding PSP pathogenesis. Overall, the efficiency and reasonable performance of OD models show great promise for tau OD.

## Acknowledgments

The authors would like to thank Kwan Wynn Tan for providing an early prototype of the data processing pipeline for OD models used in this study.

